# A role for differential gene regulation in the rapid diversification of melanic plumage coloration in the dark-eyed junco (*Junco hyemalis*)

**DOI:** 10.1101/315762

**Authors:** Mikus Abolins-Abols, Etienne Kornobis, Paolo Ribeca, Kazumasa Wakamatsu, Mark P. Peterson, Ellen D. Ketterson, Borja Milá

**Author notes:** these authors contributed equally (Mikus Abolins-Abols, Etienne Kornobis). Corresponding author: Borja Milá, Department of Biodiversity and Evolutionary Biology, National Museum of Natural Sciences, Spanish National Research Council, Calle José Gutiérrez, Abascal 2, Madrid 28006, Spain.

## Abstract

Color plays a prominent role in reproductive isolation, therefore understanding the proximal basis of pigmentation can provide insight into speciation. Subspecies of the dark-eyed junco (*Junco hyemalis*) have evolved marked differences in plumage coloration since the Last Glacial Maximum, yet whether color differences are caused by mutations in coding regions of expressed genes or are instead the result of regulatory differences remains unknown. To address this question, we studied the pigment composition and the genetic basis of coloration in two divergent subspecies, the slate-colored and Oregon juncos. We used HPLC and light microscopy to investigate pigment composition and deposition in feathers from four body areas. We then used RNAseq to compare the relative roles of differential gene expression in developing feathers and sequence divergence in transcribed loci under common garden conditions. Junco feathers differed in eumelanin and pheomelanin content and distribution. Within subspecies, in lighter feathers melanin synthesis genes were downregulated (including PMEL, TYR, TYRP1, OCA2, MLANA), ASIP was upregulated. Feathers from different body regions also showed differential expression of HOX and Wnt genes. Feathers from the same body regions that differed in color between the two subspecies showed differential expression of ASIP and three other genes (MFSD12, KCNJ13, HAND2) associated with pigmentation in other taxa. Sequence variation in the expressed genes was not related to color differences. Our findings support the hypothesis that differential regulation of a few genes can account for marked differences in coloration, a mechanism that may underlie the rapid diversification of juncos.

## INTRODUCTION

Color traits are among the most rapidly evolving phenotypes in animals and plants (Hubbard et al., 2010; Protas & Patel, 2008), and they often represent the only phenotypically diagnosable differences between species (Bourgeois et al., 2017; Campagna et al., 2016). The rapid evolution of animal color is often attributed to sexual selection (Gray & McKinnon, 2007; Lande et al., 2001; Naisbit et al., 2001), because rapidly evolving sexually selected color traits may cause prezygotic reproductive barriers due to differences in mate preference, potentially leading to reproductive isolation and speciation (Seehausen et al., 2008). Understanding the mechanisms that underlie color divergence between populations is therefore critical for a better understanding of the speciation process.

This divergence is particularly apparent in birds, where color diversity has three main components: the diversity of pigments, the patterns of pigment deposition on different parts of a feather, and the modular organization of feather tracts across the bird’s body, which may enable rapid recombination of color schemes (Badyaev, 2004, 2006). Some of the diversity in feather color has been shown to evolve as rapidly as within a few thousand years (Milá et al., 2007; Ödeen & Björklund, 2003; Zink et al., 2003), representing one of the fastest rates of evolutionary change reported in wild species. In some cases, the main genetic differences between species are in regions that encode color genes (Campagna et al., 2016; Poelstra et al., 2014), suggesting that speciation may start from only a few changes in mechanisms underlying color development.

Furthermore, specific patterns of coloration often evolve independently in distantly related species (Shapiro et al., 2013), suggesting that common mechanisms may underlie major aspects of bird color diversity by channeling color variation along specific evolutionary trajectories (Poelstra et al., 2014).

Because of the power and promise of genetic studies of color variation, the genetics of pigment production have been extensively studied in mammals and birds for the better half of the past century (Hoekstra, 2006; Hofreiter & Schöneberg, 2010; Mundy, 2005; Silvers & Russell, 1955; Yu et al., 2004). This is especially true for melanic color diversity, which has a strong genetic basis (Roulin & Ducrest, 2013). Melanic color diversity in birds is generated mainly by two pigments: eumelanin (grey, brown, black colors) and pheomelanin (yellow, red), which are produced in melanocytes, specialized pigment cells (Galván & Solano, 2016). Color differences in birds may be due to either differences in the chemical composition of melanin polymers, differential development of melanocytes, or differential distribution of melanin granules in the feather. Melanin synthesis has been shown to be regulated via numerous pathways (Hoekstra, 2006), including the melanocortin 1 receptor (MC1R) and its two ligands, the α-MSH and the Agouti signaling protein (ASIP) (Gluckman & Mundy, 2017; Yoshihara et al., 2012). Mutations in the regulatory genes, notably MC1R, have been shown to result in drastic changes in coloration in domestic and laboratory animals (Hoekstra, 2006; Kijas et al., 1998; Rieder et al., 2001; Våge et al., 1999) and, to a lesser extent, in wild species (Nachman et al., 2003; Theron et al., 2001; Uy et al., 2009).

While the role of MC1R and ASIP genes in generating color variation in some domesticated and undomesticated species is appreciated, our understanding of color evolution is nevertheless incomplete (Hoekstra, 2006). First, although sequence divergence in MC1R has been associated with melanic coloration in several cases (Theron et al., 2001; Uy et al., 2016), it often fails to explain polymorphisms (Bourgeois et al., 2016; Cheviron et al., 2006; MacDougall-Shackleton et al., 2003; Riyahi et al., 2015). Given the number of pathways that have been shown to regulate melanin production and melanocyte differentiation, this may not be surprising.

Indeed, the historical focus on a few candidate genes belies the complexity of the molecular and genetic networks that underlie melanin-based coloration (San-Jose & Roulin, 2017). Color variation in the wild can be more subtle and is often continuous, indicating complex interactions between the mechanisms that regulate local melanin production, polymerization, melanosome maturation, and deposition in the developing barbs and barbules (Arai et al., 2017; Bourgeois et al., 2017; Poelstra et al., 2014; Yang et al., 2017). To better understand the genetic basis of the natural diversity of coloration, we therefore need to expand our scope to identify additional candidate genes and mechanisms that generate color variation.

Second, we know little about the relative roles of gene expression and point mutations in coding regions in affecting color variation in the wild (Roulin & Ducrest, 2013). Although some work has been done in domesticated birds (Cooke et al., 2017; San-Jose et al., 2017), experimental work under controlled conditions aimed at understanding the role of gene regulation in affecting melanic coloration has been scarce (Ekblom et al., 2012).

In this study, we addressed both of these issues by studying gene expression underlying plumage color divergence in two plumage forms of the dark-eyed junco (*Junco hyemalis*) in a common garden environment. The dark-eyed junco complex is a quintessential example of rapid evolution of plumage color (Milá et al., 2007) and consists of at least six distinct, geographically structured subspecific forms with strikingly different plumage coloration (Nolan et al., 2002).

Recent molecular evidence indicates that the diversification within the dark-eyed junco species complex has occurred within the last 10,000 years following their post-glacial expansion in North America (Friis et al., 2016; Milá et al., 2007). Color is the main phenotypic difference between these taxa, which are otherwise morphologically similar and do not differ in their song (Nolan et al., 2002). The main color differences between junco subspecies occur on their heads, backs, and flanks. Importantly, variation in plumage traits that delineate subspecies (color of the head, amount of white on tail feathers), has also been shown to have social significance (Hill et al., 1999; Holberton et al., 1989). This suggests that the differences in junco feather color are involved in mate choice and thus may play a role in the development of assortative mating and prezygotic isolation.

We investigated the mechanisms responsible for color divergence in two forms of the dark-eyed junco: the slate-colored junco and the Oregon junco, which occur in temperate areas of Eastern and Western North America, respectively (Nolan et al., 2002). Slate-colored juncos have uniformly slate-gray upper parts, lighter gray flanks, and ventral areas (bellies), whereas Oregon juncos have black heads, brown backs, light brown flanks, and white ventral areas (Figure 1). We first asked if the two subspecies differed in the concentration of eumelanin and pheomelanin in their head, back, flank, and ventral feathers using high performance liquid chromatography (HPLC). We predicted to find more pheomelanin in the light brown-colored flanks and brown backs of Oregon juncos, compared to the gray feathers of slate-colored juncos. We then investigated if the two subspecies differed in the patterns of eumelanin and pheomelanin deposition in different feather regions (rachis, barbs and barbules) using light microscopy. To investigate the mechanisms underlying color divergence, we used RNA sequencing to characterize gene expression differences and sequence variation associated with variation in feather color between junco subspecies as well as across different body parts (head, flank, back, belly) within subspecies. Our objectives were to determine (i) whether candidate genes well known to regulate melanic coloration in domestic and some wild birds were also involved in the color differences between the two junco subspecies as well as among different feather types within each subspecies (ii) whether novel genes may be involved in this radiation that have not been known to control color in birds, and (iii) whether color differences among subspecies may be explained by point mutations in coding regions of the expressed genes or are instead the result of regulatory differences of these genes.

**Figure 1.**
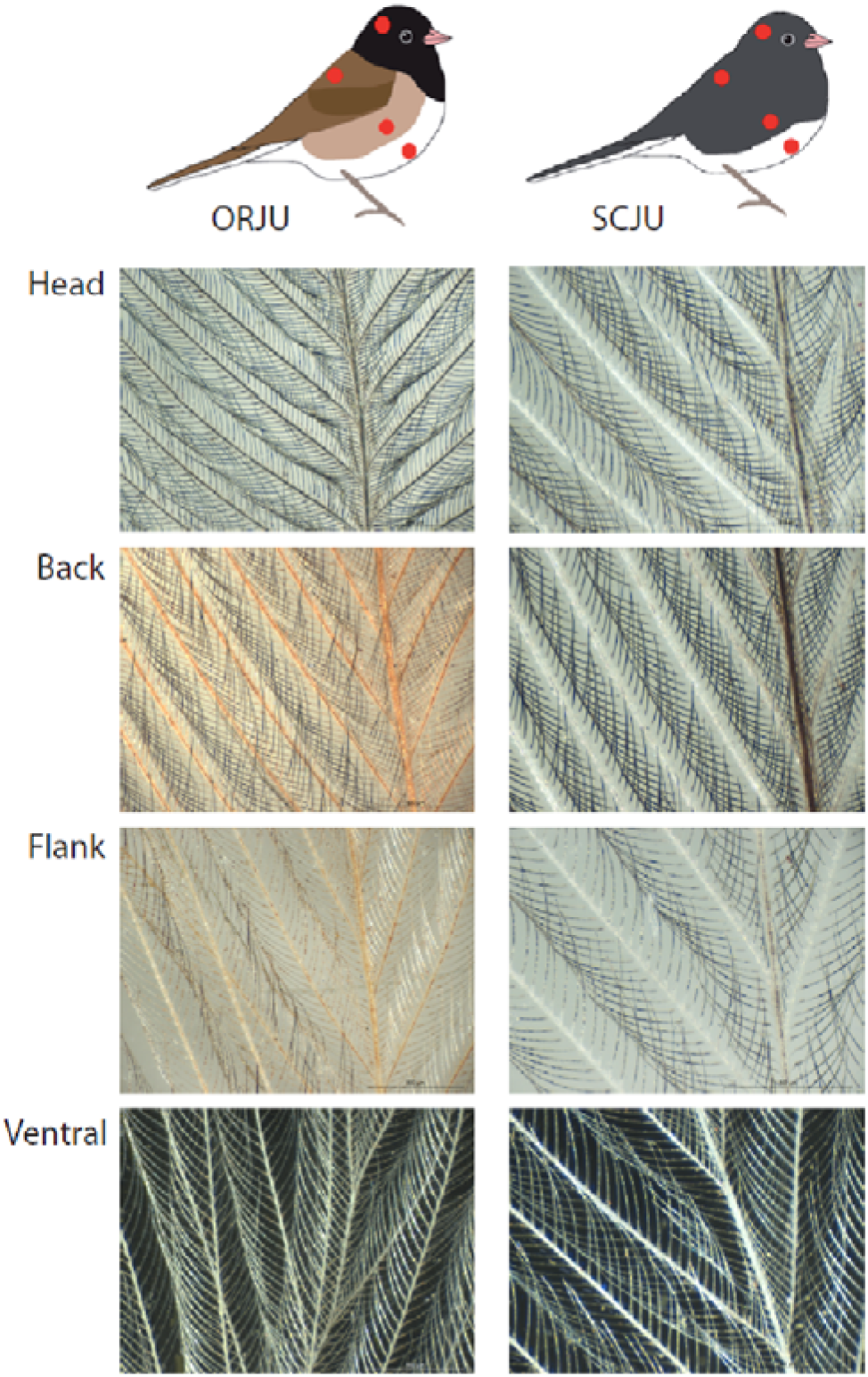
Light microscopy of junco feathers showing differential distribution of pheomelanin and eumelanin in barbules, barbs, and rachis (100x magnification). ORJU: Oregon junco; SCJU: slate-colored junco.

## MATERIALS AND METHODS

### Feather sampling and experimental design

We sampled feathers from two dark-eyed junco (*Junco hyemalis*) subspecies: the Oregon junco (*J. h. thurberi*, n=4) and the slate-colored junco (*J. h. carolinensis*, n=4). Oregon juncos (*herein abbreviated* ORJUs) were originally captured on University of California San Diego campus and the Laguna Mountains in the San Diego County, California, USA. Slate-colored juncos (*herein abbreviated* SCJUs) were captured at Mountain Lake Biological Station, Giles County, Virginia, USA. In order to reduce the influence of external factors on gene expression, birds were kept in a common-garden environment for at least 7 months (including the fall molt) prior to the collection of feathers. All birds used in the experiment were placed in cages in a single room, under identical light and temperature conditions, and were fed the same food. For each individual, we plucked mature feathers from four distinct body areas: head (coronal region of the capital feather tract), back (interscapular region of the spinal tract), flank (dorsal side of the sternal region of the ventral tract), and ventral area (belly; the ventral side of the sternal region of the ventral feather tract). For each tract, we plucked feathers from an area of 1 cm^2^. Mature feathers were used for the pigment composition and distribution analysis, while the plucking served to induce feather development in the plucked area. We monitored feather regrowth every two days following plucking. We collected developing feathers during the development of the pennaceous vane, when the first mature barbs started to erupt from the tip of the follicle (8 to 19 days following the plucking, median 11 days). To collect feather follicles, we applied a topical anesthetic to the skin, and gently plucked individual developing feathers using forceps. Six follicles were collected from each area. Plucked follicles were immediately placed on pulverized dry ice, and thereafter frozen at -80 °C until RNA extraction.

### Quantification of melanins in mature feathers

We examined the patterns of pigment deposition in the feather rachi, barbs, and barbules using a Leica MZ16A stereomicroscope at a magnification of 100X, and photographed each feather using a Leica DFC550 camera. We also quantified melanin content using high performance liquid chromatography (HPLC) to measure degradation products of pheomelanin (4-amino-3- hydroxyphenylalanine, 4-AHP, and thiazole-2,4,5-tricarboxylic acid, TTCA) and eumelanin (pyrrole-2,3,5-tricarboxylic acid, PTCA). Feather samples were homogenized with a Ten-Broeck homogenizer at a concentration of 1 mg/mL H_2_O. 100 μL (0.1 mg) aliquots were subjected to alkaline hydrogen peroxide oxidation (Ito et al., 2011) and hydroiodic acid hydrolysis (Wakamatsu et al., 2002).

### RNA extraction and cDNA library preparation and sequencing

To analyze gene expression, we created 32 separate libraries – one for each of the four body areas (head, back, flank, ventral) for each of the eight individuals (four per morph). We used 6 developing feathers for each library to ensure sufficient RNA recovery. RNA was extracted in TRIzol following manufacturer directions (Invitrogen, Carlsbad, CA, USA). cDNA libraries were prepared for the polyA-enriched fraction of the transcriptome at Macrogen Inc., South Korea, using Illumina Truseq RNA technology and sequenced in an Illumina HiSeq2000 platform. Only 3 of the ventral region libraries were sequenced per subspecies and one SCJU-Back library failed, so that a total of 29 libraries were successfully sequenced (Oregon junco: 4 × Back, 4 × Flank, 4 × Head, and 3 × Ventral; slate-colored junco: 3 × Back, 4 × Flank, 4 × Head, and 3 × Ventral). The sequencing runs produced a total of 129 Gb of cDNA, representing an average of 22 million high quality read pairs (each about 100 bp long, more than 94% of the bases with a quality above/below or equal to Q20) per library. The raw read datasets are available at ArrayExpress database at EMBL-EBI (www.ebi.ac.uk/arrayexpress) under accession number E-MTAB-6794.

### Mapping of reads

Spliced mapping was performed against the closest and most complete reference genome available, the zebra finch (*Taeniopygia guttata*, v3.2.4) genome with annotation v3.2.4.89. In order to map and obtain the read counts per transcript we used the Gemtools RNA-sequencing pipeline version 1.6, which is based on the GEM mapper (Marco-Sola et al., 2012) and is an update of the workflow used in (Lappalainen et al., 2013). Because the overall quality of the reads observed with FastQC was high and because the quality of the reads is taken into account during the mapping with GEM, no preliminary read cleaning was performed. Mapping statistics were computed with Gemtools and SAMTOOLS 1.2 *flagstat* (Li et al., 2009).

### Differential regulation analysis

Expression quantification was performed at the gene level, using FeatureCounts (Liao et al., 2014) and the genome annotation v3.2.4.89. Differential regulation analysis was conducted using the edgeR package in R (Robinson et al., 2010) with the read counts at the gene level and comparing expression in body parts within and among subspecies. Normalized read Counts Per Million (CPM) were calculated using the TMM method (Robinson & Oshlack, 2010). Differential regulation was tested using a generalized linear model approach (FDR <0.05) comparing each tissue against each other (Table 2). For further analysis of the differentially expressed genes, we focused on those that were associated with GO term “pigmentation” (GO:0043473). Genes and GO terms were matched with BioMart filtering API in Ensembl (Cunningham et al., 2014). This led to a list of 59 genes related to pigmentation to which we added the MLANA gene which was absent from the GO query results. GO term enrichment analyses were performed with the R package topGO (Alexa & Rahnenfuhrer, 2016) for differentially expressed genes in pairwise comparisons between feather types (within and between subspecies), using Fisher’s test to calculate the significance of gene enrichment. Only terms that had more than 5 annotated genes and included more than 2 significantly expressed genes are reported.

### Variant calling

Read mappings from the GEM output were further processed for variant (single nucleotide polymorphisms (SNPs) and insertions/deletions) calling in the transcribed genes. Read groups and duplicate markings were added to the bam files using the AddOrReplaceReadGroups and MarkDuplicates commands from the PICARD package (http://broadinstitute.github.io/picard). We then used GATK (McKenna et al., 2010) to identify putative SNPs and indels. We followed the guidelines from the GATK best practices for variant calling from RNA-seq data (https://www.broadinstitute.org/gatk/guide) and from (De Wit et al., 2015). Variant calls were filtered using GATK (filters used: FS > 30.0; QD < 2.0, window 35, cluster 3) and VCFtools ((Danecek et al., 2011), --max-missing 0.25, --mac 1, --min-alleles 2, --minDP 6, --minGQ 10).

Variant locations on the genome were identified using in-house python scripts. Weir and Cockerham *F*_ST_ values between the two forms ORJU and SCJU were computed using VCFtools. A GO term enrichment analysis was performed on the genes showing variants with *F*_ST_ values equal to one.

## RESULTS

### Distribution and quantification of pigments

Inspection of mature feathers from various body parts of SCJU and ORJU individuals with light microscopy color differences between subspecies are in part to do differential pigmentation of rachi, barbs and barbules (Figure 1). In the black feathers from ORJU heads, rachis, barbs, and barbules were uniformly darkly colored, suggesting predominance of eumelanin pigmentation. In contrast, in ORJU back and flank feathers, only barbules showed dark coloration consistent with eumelanin, while barbs and rachi showed orange-brown coloration, consistent with pheomelanin predominance. Feathers from the gray heads, backs, and flanks of SCJUs had darkly pigmented rachi and barbules, yet barbs contained no apparent pigment. The pennaceous part of white ventral feathers from both subspecies showed no apparent pigmentation.

Eumelanin and pheomelanin concentrations quantified using HPLC were overall consistent with the light microscopy observations. Eumelanins were found in feathers from both subspecies and all body parts, whereas pheomelanins were absent in all SCJU feather samples, but present in ORJU back, flank feathers, as well as in the black ORJU heads where it may be masked by eumelanin (Table 1). ORJU back feathers (brown) showed the highest concentration of pheomelanin. We also found eumelanin in the white ventral feathers. In these feathers, the visible distal vane of the ventral feathers is white, while the more proximal feather plumes, hidden by other vanes, are gray.

**Table 1.** Degradation products of eumelanin and pheomelanin pigments measured by HPLC in feathers from different body parts of Oregon and slate-colored juncos. Values (n = 4) represent sample means and standard deviations (in parentheses).

|  | Oregon junco |  |  | Slate-colored junco |  |  |
| --- | --- | --- | --- | --- | --- | --- |
|  | Eumelanin<br>(ng/mg) | Pheomelanin<br>(ng/mg) |  | Eumelanin<br>(ng/mg) | Pheomelanin<br>(ng/mg) |  |
|  | PTCA | 4-AHP | TTCA | PTCA | 4-AHP | TTCA |
| Head | 2468 (332) | 46 (8) | 207 (117) | 1967 (314) | <9 | <94 |
| Back | 1075 (189) | 165 (38) | 369 (28) | 1796 (163) | <9 | <94 |
| Flank | 2195 (764) | 46 (13) | 148 (24) | 2707 (323) | <9 | <94 |
| Ventral | 2063 (535) | 36 (7) | 132 (100) | 2156 (408) | <9 | <94 |

### Differential gene expression between feather types within subspecies

Using the Gemtools pipeline, an average of 87.1% (min: 83.6%, max: 90.4%) of all reads were mapped to the zebra finch genome and an average of 61% (min: 59.1%, max: 63.5%) were properly paired. The relatively low figures are not surprising given the evolutionary distance between junco and zebra finch, which is the closest species for which a high-quality reference genome exists. Overall, 346 genes were differentially expressed at a statistically significant level (FDR threshold of 0.05) between different feather tracts within subspecies (Table 2, Table S1). Of these, 304 were differentially expressed in Oregon juncos, and 112 were differentially expressed in slate-colored juncos (overlap of 70 genes).

**Table 2.** Number of differentially regulated genes for each comparison between junco forms and body parts (two-fold change, down-regulated/up-regulated). Comparisons of the same tissue between sub-species is highlighted in grey. ORJU: Oregon junco; SCJU: Slate-colored junco; B: back; F: flank; V: ventral; H: head.

|  | ORJU-B | ORJU-F | ORJU-V | SCJU-H | SCJU-B | SCJU-F | SCJU-V |
| --- | --- | --- | --- | --- | --- | --- | --- |
| ORJU-H | 49/17 | 65/31 | 119/79 | 6/1 | 67/12 | 36/8 | 201/114 |
| ORJU-B |  | 3/3 | 22/46 | 10/28 | 1/0 | 8/6 | 38/37 |
| ORJU-F |  |  | 36/41 | 13/16 | 24/11 | 1/3 | 67/44 |
| ORJU-V |  |  |  | 61/77 | 71/25 | 45/57 | 0/0 |
| SCJU-H |  |  |  |  | 4/0 | 0/0 | 42/32 |
| SCJU-B |  |  |  |  |  | 1/2 | 8/16 |
| SCJU-F |  |  |  |  |  |  | 29/17 |

Among the significantly differentially expressed genes between body parts in ORJU were several members of the canonical melanin synthesis pathway –TYRP1, TYR, OCA2, RAB38, SLC45A2, SLC24A5, PMEL, and MLANA (Figure 2). Most of these genes were downregulated in developing white ventral feathers compared to colored feathers (Figure 3). Within SCJU, the qualitative patterns of expression of these genes were similar, although only SLC45A2 was significantly downregulated in SCJU ventral feathers compared to other body regions.

**Figure 2.**
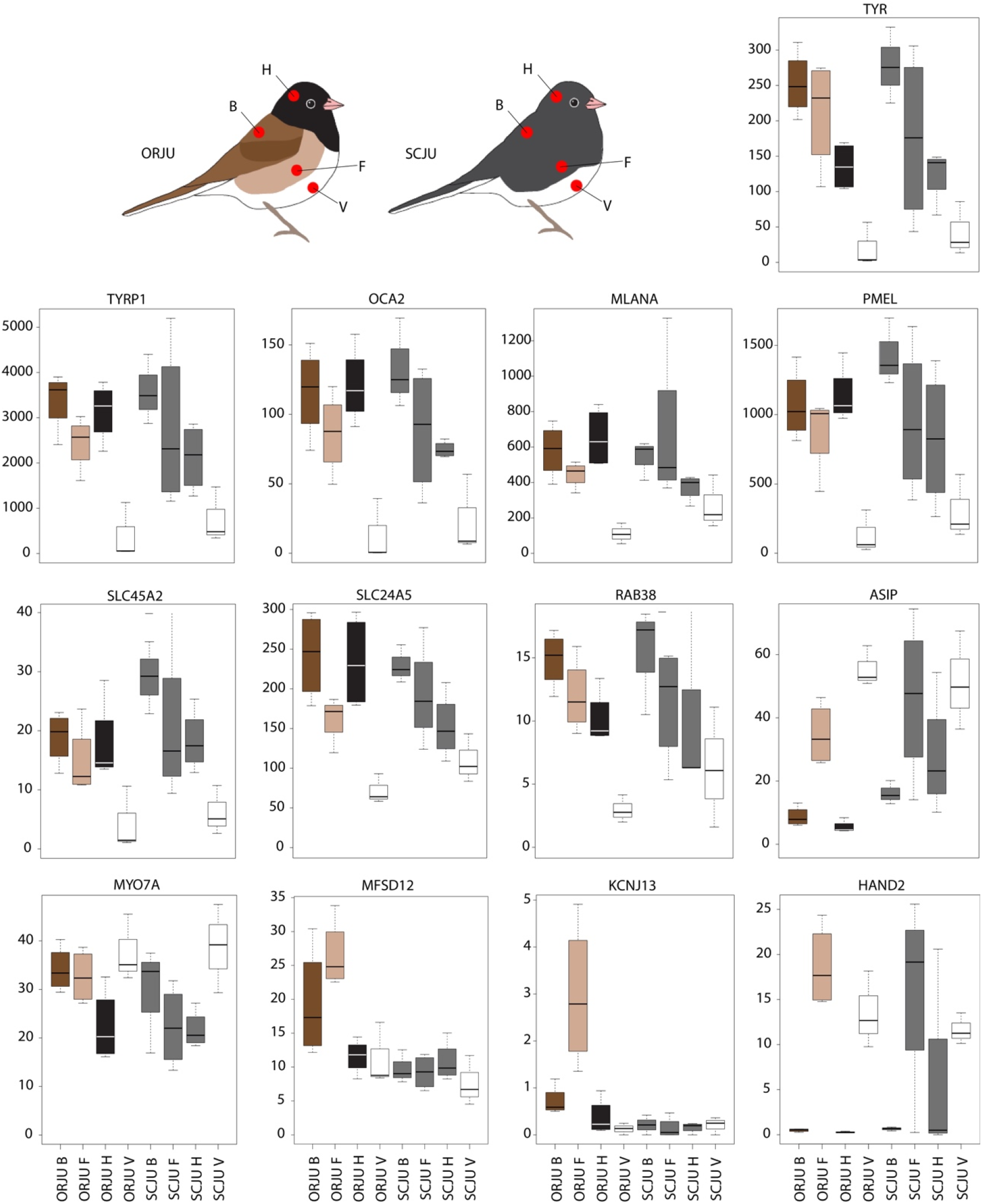
Expression (normalized CPMs) of significantly differentially regulated genes associated with color development between subspecies and body regions. ORJU: Oregon junco; SCJU: slate-colored junco; H: head; B: back; F: flank; V: ventral.

**Figure 3.**
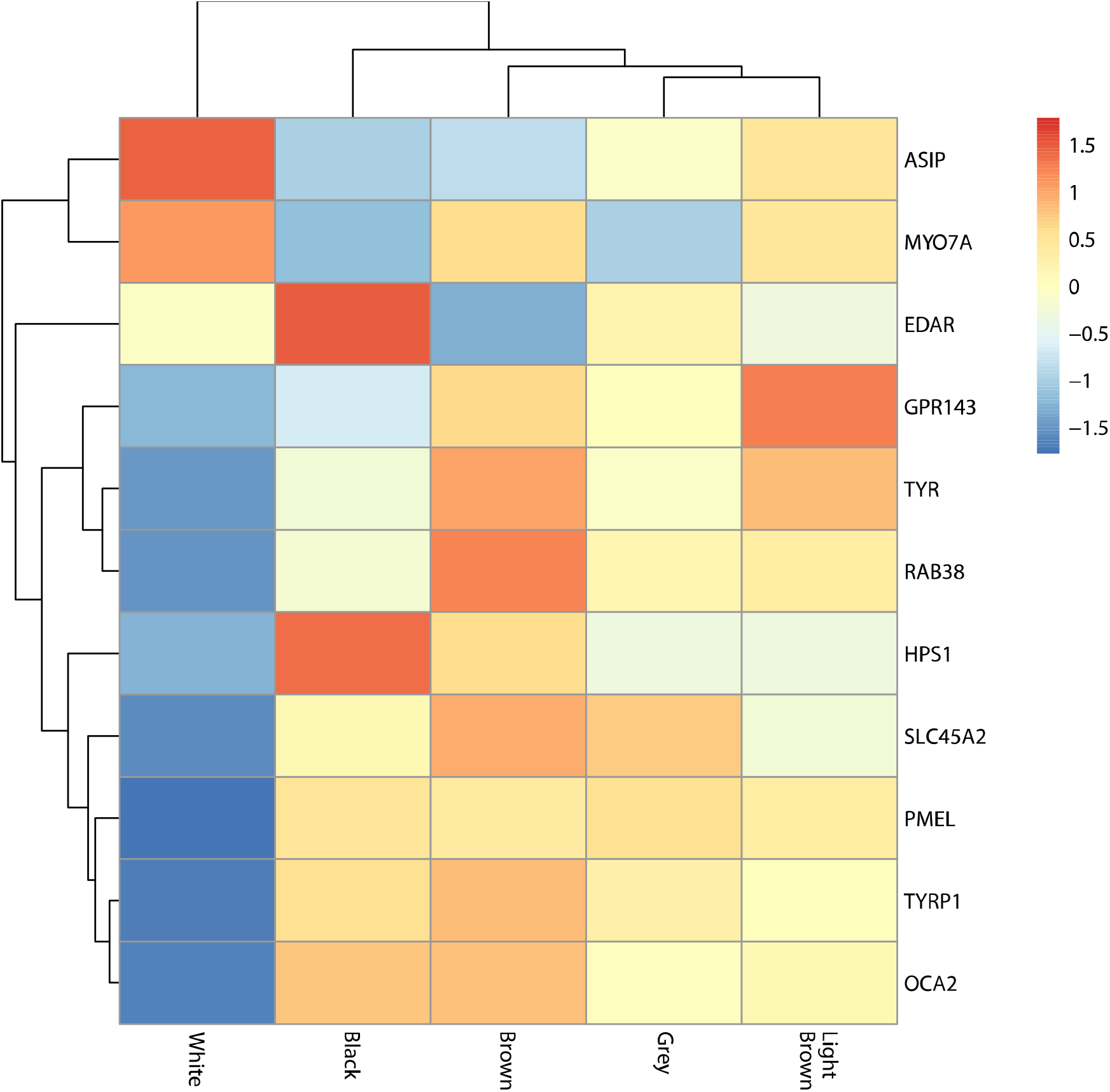
Heat map based on the median of counts (normalized CPMs) feather colors for the 11 pigment-associated genes differentially regulated in any comparison between colored feathers against white (ventral) feathers.

In ORJU, the white ventral feathers expressed significantly more ASIP, an inhibitor of eumelanin synthesis, compared to head and back feathers. In both ORJU and SCJU, feathers also showed differential regulation of Wnt signaling pathway components, including SFRP1 and DKK3, both Wnt signaling inhibitors (Figure 4). DKK3 expression was lower in the black head feathers compared to the lighter back, flank, and ventral feathers in both ORJU and SCJU, while SFRP1 expression was lower in the dark ORJU feathers compared to their white ventral feathers. On the other hand, another Wnt-signaling pathway gene, FRZB, was upregulated in SCJU head and back feathers compared to the white ventral feathers (Figure 4).

**Figure 4.**
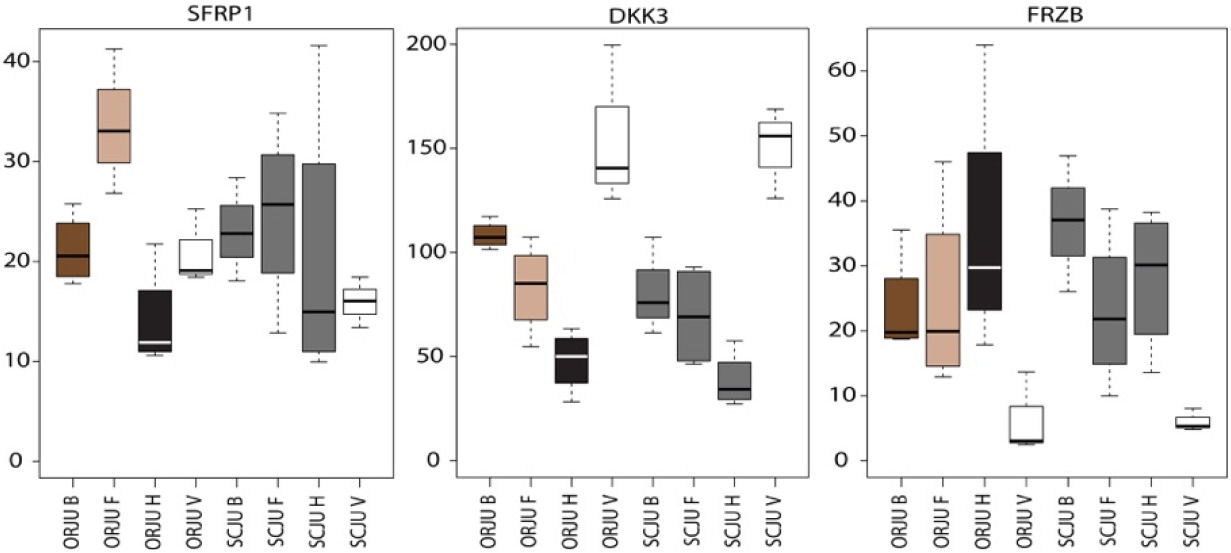
Expression (normalized CPMs) of significantly differentially regulated genes associated with Wnt signaling between subspecies and body regions. ORJU: Oregon junco; SCJU: slate-colored junco; H: head; B: back; F: flank; V: ventral.

Among other significantly differentially expressed genes between the different feather types were members of the HOX gene group. In ORJU, ten HOX genes were downregulated in the head feathers compared to back, flank, and ventral feathers. In SCJU, only two HOX genes were differentially regulated between feather types (Figure 5). Some HOX genes were differentially expressed with respect to body region rather than melanin type. For example, HOXA2 and HOXB7 were up and down-regulated, respectively, in the head feathers of both subspecies, whereas HOXB8 was upregulated only in developing ventral feathers.

**Figure 5.**
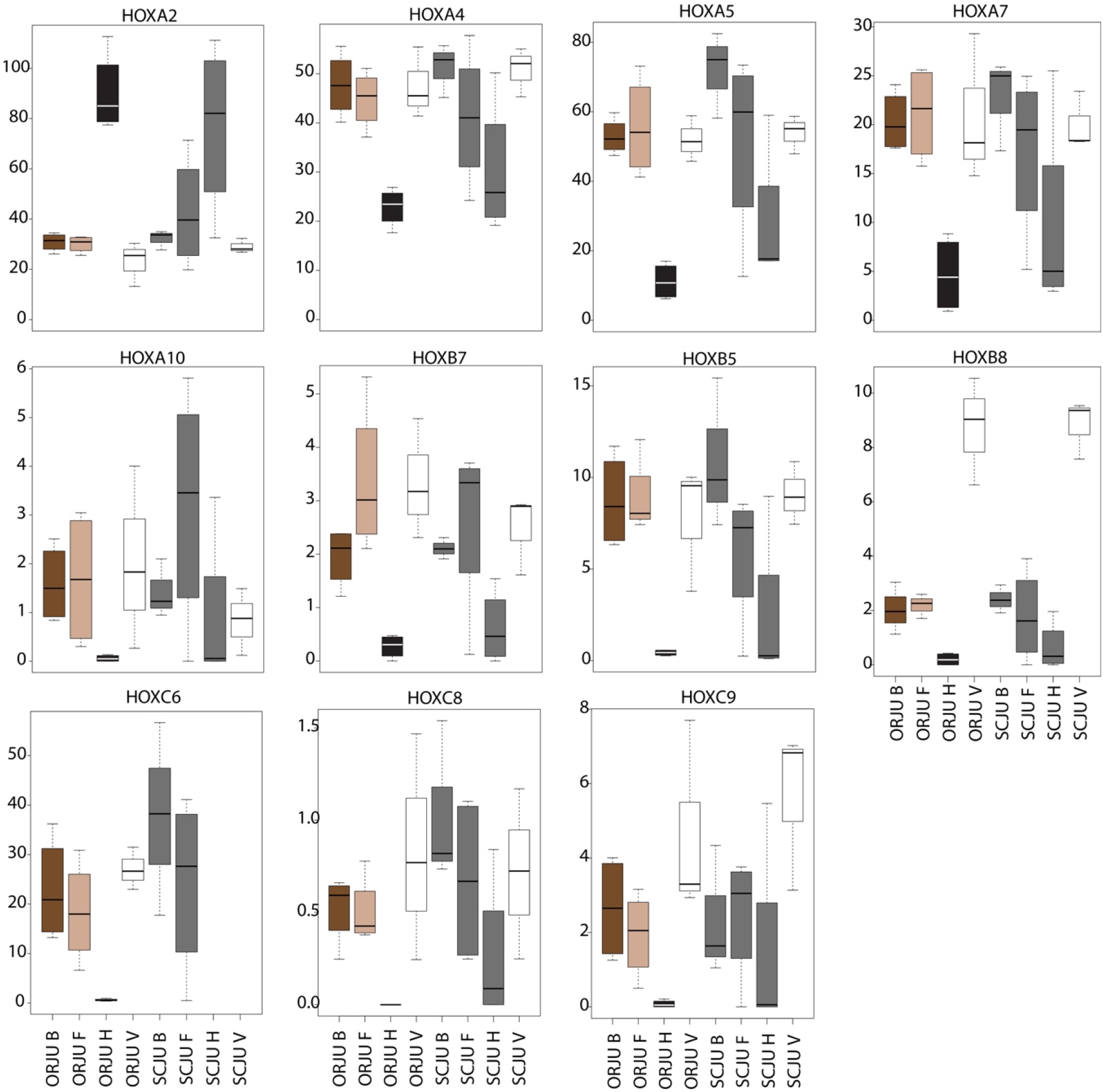
Expression (normalized CPMs) of significantly differentially expressed HOX genes between subspecies and body regions. ORJU: Oregon junco; SCJU: slate-colored junco; H: head; B: back; F: flank; V: ventral.

GO categories that were significantly enriched among the differentially regulated genes between feather types included categories related to pigmentation (e.g. melanin biosynthetic process, pigment granule organization, pigment cell differentiation; all significantly enriched between ventral feathers and colored feathers in ORJUs) as well as morphogenesis (e.g. developmental process, appendage development, tissue morphogenesis, all between head feathers and other body feathers in both ORJUs and SCJUs)(Table S2).

### Differential gene expression between subspecies

Only 10 genes (Table S3) were significantly differentially regulated between the same feather tracts across the two subspecies (Table 2). Of these, ASIP has been linked to pigment variation in birds, and three other genes (MFSD12, KCNJ13, and HAND2) have been associated with pigment production in other vertebrates. ASIP and HAND2 were more highly expressed in the gray SCJU heads compared to black ORJU heads, while MFSD12 and KCNJ13 were more highly expressed in the light brown ORJU flanks compared to the grey SCJU flanks (Figure 2). Most of the differential expression (7 out of 10 genes) was between developing ORJU and SCJU head feathers (black vs gray). Only one gene (FAM172A) was significantly differentially expressed between all three colored feather tract comparisons.

### Sequence variation between subspecies

A total of 57,214 variant sites were identified between the two morphs. Out of these, only 43 variant sites (located in 20 different genes) were segregating between the two morphs with *F*_ST_=1, but none were located in the pigmentation related gene list obtained from Ensembl. The highest *F*_ST_ value observed for a pigmentation related gene was 0.55 at the FIG4 gene.

Interestingly, the group of 20 genes with a fixed SNP contained two genes associated to the Wnt signaling pathway, FZD4 and APC.

## DISCUSSION

Color variation between the dark-eyed junco subspecies represents one of the best examples of rapid plumage color evolution in the wild. To understand the mechanisms that underlie this diversity, we characterized the pigment composition and deposition patterns in mature feathers and used RNAseq to ask if differences in coloration between two distinct junco subspecies are explained by differences in gene expression in developing feathers or by coding differences in the expressed genes. We show that coloration differences between subspecies are due to the differential deposition of eumelanin and pheomelanin in different parts of the birds’ feathers, and that variation among body parts within and across subspecies results from the differential regulation of a potentially small set of genes rather than from point mutations in their coding regions.

### Phenotypic difference in melanin deposition

Slate-colored and Oregon junco subspecies differed in the coloration of their flanks, backs, and heads. As proposed previously (Miller, 1941), the differences in coloration on a phenotypic level were explained by differences in the type of pigment deposited in the feathers, as well as the pattern in which this pigment was deposited in the rachi, barbs, and barbules (Figure 1). Eumelanin was found in both subspecies and all feather types. Pheomelanin was present in all body parts of black- and brown-colored ORJUs, whereas it was below the detection limit in the uniformly gray SCJU feathers (Table 1). Pheomelanin values in ORJU were higher for back, yet values for head, flank, and ventral area were very similar, even though head feathers are black, flank is light brown, and ventral area is white. Similar patterns of color differences despite similar pheomelanin levels have also been shown in human hair (Ito et al., 2011) and human skin (Bino et al., 2015). This can be explained by the casing model, which proposes that in melanosomes that contain both pigments, pheomelanin is produced first, followed by synthesis of eumelanin, which surrounds the pheomelanin core (Ito & Wakamatsu, 2008). Interestingly, pigment deposition in feathers was not uniform: the grey SCJU feathers had pigmented rachi and barbules, whereas barbs appeared unpigmented. In contrast, in ORJU, rachi and barbs of flank and back feathers showed orange pigmentation, likely due to a presence of pheomelanin, whereas barbules were much darker, indicating a predominance (or casing) of eumelanin. In ORJU heads, feather rachi, barbs, and barbules were all dark, suggesting a predominance of eumelanin pigment. These observations suggest that color differences between Oregon and slate-colored juncos are due to regulation of both melanin synthesis and the differential migration of the mature melanosomes in the developing feather matrix. Recent studies have also suggested that color variation in feathers may be a result of the ratios of different melanin moieties (chemical variants) in the feathers (Galván & Wakamatsu, 2016).

### Differences in expression between subspecies

We were able to detect only a handful of genes that were differentially regulated at a statistically significant level between the same body parts of ORJUs and SCJUs (Table 2). The grey SCJU heads expressed less ASIP and HAND2 compared to the black ORJU heads (Figure 2). ASIP, which encodes the Agouti-signaling peptide, is an inverse agonist to melanocortin-1 receptor (MC1R), one of central regulators of melanin synthesis in birds and mammals (Manceau et al., 2011; Mundy, 2005). Increased expression/signaling by ASIP has been shown to lead to increased synthesis of pheomelanin (Roulin & Ducrest, 2013) or arrest of melanin synthesis, leading to absence of pigmentation (Lin et al., 2013). Higher expression of ASIP in grey head feathers of SCJU suggests that ASIP may be lowering the production of melanin in these feathers, in contrast to the black ORJU feathers. ASIP has been shown to explain color variation in a wide variety of taxa (Martin & Orgogozo, 2013). Our study adds to this body of literature and suggests that the regulation of MC1R by ASIP can lead to rapid changes in the color hue.

HAND2 encodes a transcription factor that has been shown to be important in regulating cell fate during the development of various organs and limbs (Yelon et al., 2000). It is also an upstream regulator of an important patterning gene, sonic hedgehog (SHH) (Xiong et al., 2009). HAND2 has been shown to be differentially expressed in cichlid fish fins that differ in color, indicating that HAND2 may be responsible for regulation of pigment cell development or function (Santos et al., 2016). Because of this, we hypothesize that HAND2, or genes downstream in the SHH pathway, may be involved in regulation of melanin production or differential deposition of mature melanosomes in the rachi, barbs, and barbules, which are distinct tissue types in the developing feather.

The light brown pheomelanin-rich ORJU flank feathers expressed more MFSD12 and KCNJ13 compared to the grey SCJU feathers (Figure 2). MFSD12 was recently identified as one of the principal genes regulating skin color in humans and coat color in mice (Crawford et al., 2017). Knockdown studies in mice have shown that MFSD12 inhibits eumelanin synthesis while being required for pheomelanin synthesis (Crawford et al., 2017). This suggests a conserved role of this gene across major vertebrate groups and calls for further study of the role of this gene in generation phenotypic variation. KCNJ12 has been associated with changes in pigmentation patterns on zebra fishes (Haffter et al., 1996). In zebra fishes, the potassium channel encoded by this gene regulates the interactions between melanophores and xantophores (Singh & Nüsslein-Volhard, 2015). KCNJ13 function is unknown in birds, but it may regulate interactions between pigment cells and the surrounding cellular matrix, perhaps being responsible for the differential pigment deposition in barbs and barbules.

### Differences in gene expression between feather types within subspecies

Compared to head, back, and flank, the white ventral ORJU feathers had lower expression of genes that encode four main regulators/catalysts of melanin production in the melanosome: TYR (tyrosinase), TYRP1 (tyrosinase-related protein 1), OCA2 (OCA2 melanosomal transmembrane protein), and SLC45A2 (Solute Carrier Family 45 Member 2, Figure 2). TYR and TYRP1 catalyze reactions that lead to the conversion of tyrosine to melanin, while the role of SLC45A2 and OCA2 in bird melanocytes is less well understood (Galván & Solano, 2016). Mutations in SLC45A2 have been shown to inhibit the synthesis of pheomelanin, suggesting that it may control pheomelanin production (Gunnarsson et al., 2007). White feathers also showed lower expression of MLANA/MART1 and GPR143/OA1 which regulate melanosome biogenesis and maturation (Aydin et al., 2012; Cortese et al., 2005; Schiaffino & Tacchetti, 2005)(Figure 2).

Melanin synthesis in birds is regulated by at least three semi-independent pathways: MC1R, Wnt, and MAPK pathways (Poelstra et al., 2014). White ventral and light brown flank ORJU feathers expressed more ASIP compared to ORJU heads or backs. ASIP is an inverse agonist to MC1R, indicating that signaling along this pathway may be responsible for the suppression of melanin synthesis in the white ventral and light brown flank feathers. We also found that ventral feathers expressed significantly less SFRP1, a gene encoding frizzled-related protein that plays an important role in Wnt signaling, and significantly more DKK3, a Wnt-signaling inhibitor (Figure 4). Genes from the DKK family have been shown to suppress melanocyte function and proliferation (Yamaguchi et al., 2007). This suggests that white feather color may also result from inhibition of Wnt-activated melanin synthesis. Surprisingly, another Wnt-signaling inhibitor that is linked to melanocyte function FRZB (Thomas & Erickson, 2008), showed the opposite pattern, being expressed at lower levels in the white ventral feathers (Figure 4). This suggests that either the role of FRZB in avian melanocytes may be different compared to mammalian systems, that FRZB may be responsible for processes other than feather color (see below), or that FRZB may be involved in arresting melanocyte function following active melanin synthesis. FRZB has shown to be associated with darker pigmentation in other bird species as well, suggesting a similar function of FRZB across avian taxa (Poelstra et al., 2015).

### Differential expression of genes associated with feather type

It is important to note that, instead of regulating color, many of the differentially expressed genes between different feather types may be regulating feather morphology. Alternatively, these differences may reflect differences in developmental timing, as we could not ensure that feathers from different body regions were collected at the exact same developmental stage. Poelstra et al. (2015) differentiated between the putative functions (color vs. shape) of differentially expressed genes by asking if expression differences between feather types persist across taxa, given that at least in one taxon the color is the same between feather types. Because SCJUs have grey feathers on their heads, backs, and flanks, we applied this logic to ask if genes differentially expressed in these feathers in SCJU were also differentially expressed in the equivalent comparisons in ORJU. We found only three genes that were consistently differentially expressed between feather types across subspecies. Only one of these genes (IL17REL) was annotated (lower expression in back compared to flank in both subspecies), but it has not been linked to feather development before.

### Differential expression of HOX and Wnt genes

We found differential expression in 11 HOX genes between white and darker feathers, although our experimental design does not allow us to assign precise functions to these genes (Komiya & Habas, 2008)(Figure 5). HOX genes are transcription factors that regulate morphogenesis via their time- and location-specific expression (Krumlauf, 1994). All but one (HOXA2) showed lower expression in the black ORJU head feathers compared to the ORJU flank, back, and belly feathers. Among these, HOXC8 has been shown to regulate feather morphology in chickens, and its misexpression can turn head feathers into body-like feathers (Boer et al., 2017; Wang et al., 2012). SCJU head feathers showed qualitatively similar HOX expression patterns as ORJU head feathers, although only two HOX genes (HOXA2, HOXB8) were significantly differentially expressed between SCJU head and ventral feathers. These qualitatively similar patterns suggests that HOX genes may be either regulating head-specific feather morphology (head feathers are much smaller) or reflect differences in the developmental stage of feathers across different body regions at the time of tissue collection. On the other hand, the strong difference in HOX expression between ORJU flank, back, and ventral feathers (which differ in pigment deposition patterns), and the near absence of such differences in the SCJU feathers (which have similar pigment deposition patterns), suggests that HOX genes may also be involved in regulation of feather color, as shown in Drosophila (Jeong et al., 2006), perhaps through their capacity to regulate cell migration (Stoll & Kroll, 2012).

In addition to HOX genes, another important signaling pathway for morphogenesis is the Wnt signaling pathway, which regulates cell fate, migration, and tissue patterning (Komiya & Habas, 2008). Therefore, differences in expression of Wnt-related genes in white and dark feathers may reflect the role of these genes in regulating feather growth or differences in feather shape.

### Sequence variation and population differentiation

We identified only 43 variant sites segregating SCJU and ORJU forms (FST=1), none of them in genes closely related to pigmentation, indicating that differential color pigmentation in the two forms are more likely due to regulatory mechanisms than to sequence variation in the coding regions of known pigmentation-related genes. Nonetheless, although weakly supported in terms of number of good quality genotypes called, segregating variants were detected in Wnt-pathway related genes, providing further support for the potential role of Wnt signaling in feather color regulation. Higher coverage sequencing as well as SNP data in non-coding introns and *cis* and *trans* regulatory sites could shed light on the implication of sequence variation in the regulation of pigmentation of the two forms. An additional possibility is that differential expression may be due to copy number variation of the underlying genes, or due to environment-driven differences in the epigenetic regulatory mechanisms. However, our common garden approach should have at least partly eliminated the possibility of environmentally-induced plumage variation.

### Rapid evolutionary change

We have shown that rapid evolution of feather color in the genus junco can be explained by changes in pigment composition and pigment distribution. While the role of pigment composition in the evolution of color is appreciated, few studies have investigated how differential pigment distribution on rachis, barbs, and barbules, contributes to color divergence (Galván, 2011). Here we demonstrate striking differences in pigment distribution in feathers between closely related subspecies. Because developing barbs and barbules occupy distinct locations in the developing feather (barbules are more peripheral in the cross-section of a developing follicle (Yu et al., 2004)), these differences could be achieved by relatively few changes in the regulation of melanosome deposition.

Indeed, our data show that feather color differences at the phenotypic level could be achieved by simple changes in gene expression involving canonical melanocyte signaling pathways and, possibly, genes that regulate pigment distribution within feathers. These findings demonstrate that drastic changes in plumage color can evolve rapidly and readily. Furthermore, although our study might have failed to identify junco-specific pigmentation genes due to the lack of a complete genomic reference for the species, our findings are consistent with the hypothesis that evolution of coloration in birds and other vertebrates involves the same molecular pathways and genes (Martin & Orgogozo, 2013). For example, ASIP has been shown to regulate color in both mammals (Manceau et al., 2011; Rieder et al., 2001; Steiner et al., 2007) and birds (Campagna et al., 2016; Lin et al., 2013; Nadeau et al., 2008). Many of the studies investigating the role of ASIP capitalize on color polymorphisms in domesticated or model organisms. Our study joins the small but growing number of studies showing that ASIP may regulate evolution of feather color in wild populations. The absence of segregating SNPs in the coding sequence of ASIP and other genes involved in pigmentation suggests that variation in the feather color between junco subspecies may be due to variation in the regulatory regions of these genes, or in the coding or regulatory sequences of upstream transcription factors.

In addition to ASIP, we identify three other candidate genes - MFSD12, KCNJ13, and HAND2 – that have been shown to be important in vertebrate color development and evolution but have not, to our knowledge, been linked to color differences in birds. Identification of such genes is important because major candidate genes for feather color, such as ASIP, explain only part of the phenotypic diversity observed in the wild (San-Jose & Roulin, 2017). Future studies should further investigate the precise role that these genes may play in regulating melanin synthesis or melanosome distribution.

## ACKNOWLEDGEMENTS

Funding was provided by a grant from the Spanish Ministry of Science and Innovation (CGL-2011-25866) to BM.

## AUTHOR CONTRIBUTIONS

B.M. and E.D.K designed the study. M.A.A. and M.P.P. conducted the common-garden experiments. K.W. analyzed the chemical composition of feathers. E.K., P.R. and M.A.A. analyzed the transcriptomic data. M.A.A., E.K., and B.M. wrote the paper with input from all authors. B.M. provided funding for the project.

## DATA ACCESSIBILITY STATEMENT

RNA-seq data have been deposited in the ArrayExpress database at EMBL-EBI

(www.ebi.ac.uk/arrayexpress) under accession number E-MTAB-6794.

